# Unveiling value functions in social cognition with multi-agent inverse reinforcement learning

**DOI:** 10.1101/2024.10.09.617461

**Authors:** Yusi Chen, Yuan Cheng, Myungji Kwak, Angela Radulescu, Herbert Zheng Wu

## Abstract

Social behavior requires individuals to consider not only their own goals but also those of others. Latent reward functions that encode such goals can be recovered from behavior using inverse reinforcement learning in single-agent settings. However, extending this to multi-agent interactions is challenging, as joint reward functions are defined over joint state spaces that grow exponentially with the number of agents. Existing approaches address this complexity by imposing restrictive assumptions on social interactions, limiting generality and interpretability. Here we show that joint reward functions governing social interactions can be efficiently represented through decomposition into individual reward maps for each agent and low-dimensional interaction terms. Building on this formulation, we introduce a general multi-agent inverse reinforcement learning framework (MAIRL) that enables scalable and interpretable inference of latent reward representations directly from behavior. In simulated social tasks with known ground-truth rewards, MAIRL recovered the rewarded states and imitated expert behavior as accurately as or better than existing multi-agent inverse reinforcement learning methods. Applying MAIRL to mouse and primate social tasks, we recover structured reward maps that reflect distinct social roles and interaction strategies during group behavior. These results establish MAIRL as a general framework for identifying latent objectives underlying multi-agent behavior across species and paradigms.

## 1 Introduction

Social interactions are central to survival and daily life [1–5], yet deciphering the computations that govern them remains a fundamental challenge. In these settings, one’s decisions both influence and depend on the actions of others, making it difficult to determine how the brain evaluates outcomes and integrates social information. Inverse reinforcement learning (IRL) is a powerful approach for studying single-agent behavior by inferring its latent reward functions that encode an agent’s goals [6–9]. This framework has been instrumental in linking behavior to neural representations [10, 11] and in understanding how these computations break down in neuropsychiatric disorders [12, 13]. Extending IRL to social interactions is therefore essential for uncovering the computational principles underlying social cognition and its dysfunction.

However, applying this framework to social interactions introduces key challenges: the latent reward functions are defined over the joint state space of interacting agents, which grows exponentially with the number of agents. To manage this complexity, many existing cognitive and neuroscience models rely on strong structural simplifications, limiting their applicability across diverse social tasks. One common strategy embeds social influences as latent variables within a single-agent partially observable Markov decision process (POMDP) framework [14–17]. While effective for capturing behavior, these approaches obscure the structure of social interactions by collapsing them into unobserved latent factors. Other approaches [18, 19] adopted multi-agent Markov Decision Processes (MDPs) to infer simplified, often binary, social intentions to improve task performance, but do not recover the full reward structure. Related work [20] assumes distinct self-reward functions across cognitive hierarchies, conflating task-related reward structure with agents’ recursive reasoning. These limitations highlight the need for a framework that can recover the complete and interpretable reward functions directly from multi-agent behavior. This would enable direct inference of how agents represent both the environment and their social partners, disentangling task structure from social reasoning and allowing quantitative comparison of strategies.

To enable inverse inference of latent objectives from behavior, we first construct a generative forward model that links reward functions to observable social decisions through soft value iteration under well-defined computational assumptions. We formalize social tasks within a multi-agent reinforcement learning (MARL) framework [21], which specifies how multiple agents act and interact in a shared environment. We show that joint reward functions governing social interactions can be efficiently represented through value decomposition, which factorizes them into individual reward components for each agent and low-dimensional interaction terms. This representation reduces the dimensionality of the inference problem while preserving task-relevant interactions between agents, enabling tractable and interpretable recovery of joint reward functions from behavior. We show analytically and numerically that, in typical social tasks where joint rewards are sparse, this decomposition accurately reconstructs the joint reward function, from which the corresponding action-value and state-value functions are obtained by soft value iteration. Within this framework, task structure is encoded in the joint reward function, while agents’ cognitive hierarchy is captured by recursive reasoning [22], with recursion depth corresponding to theory-of-mind capacity.

Building on this formulation, we develop a multi-agent inverse reinforcement learning (MAIRL) framework [6, 7, 23] that infers decomposed reward representations from observed behavior, thereby recovering agents’ internal representations of both the environment and their social partners. This inference algorithm accurately recovers the ground-truth reward functions in simulated social foraging tasks, validating the approach under controlled conditions. Benchmarking against two existing multi-agent IRL methods showed that the decomposition preserves policy imitation while improving reward recovery, particularly when the reward depends on the relational structure between agents. We then apply MAIRL to empirical data from mouse cooperative behavior, revealing how asymmetric value learning across agents gives rise to leadership and followership. In a non-cooperative primate task, MAIRL uncovers mutual prediction signals correlated with social hierarchy. Together, these results demonstrate that value decomposition through MAIRL enables interpretable, scalable inference of the goals and strategies underlying social interactions across tasks and species.

## 2 Results

### 2.1 MAIRL with value decomposition

Social interactions between two agents can be modeled as a two-agent Markov Decision Process (MDP) ^1^ The joint state space was denoted as *S* = *S*_1_ × *S*_2_, with joint state *s* = (*s*_1_, *s*_2_) combining two agents’ individual states, and the joint action space as *A* = *A* _1_ × *A*_2_, with joint action *a* = (*a*_1_, *a*_2_). The transition kernel *P* (*s*^*′*^| *s, a*) specifies the environment dynamics, *r*^*i*^ denotes the reward function of agent *i*, and *γ* ∈ [0, 1] is the discount factor. The reward function *r*^*i*^ assigns a scalar value to agent *i* when all agents transition from joint state *s*∈ *S* to *s*^*′*^∈ *S* under joint action *a*∈ *A* . Consequently, its domain is defined over the joint state–action space, whose cardinality grows exponentially with the number of agents. When reward functions are identical across agents (*r* = *r*^1^ = *r*^2^), the formulation captures cooperative interactions; when they differ, it accommodates competitive or mixed-motive social behaviors. For clarity, we focus on the identical-interest setting and omit agent superscripts below.

In inverse reinforcement learning [6, 7, 24], given a known environment {*P, S, A, γ*} and expert trajectories 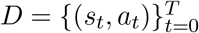, the goal is to recover a reward function *r* that maximizes the likelihood *P* (*D*|*r*). Under a conditional independence assumption, the trajectory likelihood factorizes as

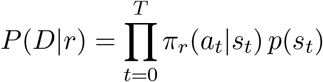

where policy *π* is induced by the underlying value functions and thus implicitly parameterized by reward *r*. Details of the problem formulation are provided in Sections 4.1, 4.3 and 4.4, where a probabilistic generative model for the joint policy *π*(*a* | *s*) is formally defined by combining a multi-agent reinforcement learning framework with maximum-entropy value iteration and recursive reasoning.

A key challenge is that joint reward functions in multi-agent settings are defined over a space that grows exponentially with the number of agents, making them difficult to compute and interpret. Here we show that these functions can be effectively represented through **value decomposition**, which expresses the joint reward as individual components of each agent and low-dimensional interaction terms:

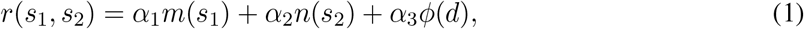

where *m* and *n* denote reward maps over individual states, *ϕ*(*d*) is an interaction term depending on the behavior (e.g., Dijkstra distance *d*, defined as number of minimal steps), and *α*_*k*_ are scalar weights. This decomposition reduces the parameter space from |*S*_1_| |*S*_2_| to| *S*_1_| + |*S*_2_| + |*d*|, while preserving the task-relevant structure of social interactions. Although we refer to this formulation as value decomposition following standard terminology, the quantity inferred by MAIRL is the reward function *r*. Action-value and state-value functions *Q* and *V* are instead obtained by soft value iteration on the inferred reward (Eq. 7); because this operation involves a log-sum-exp over actions and successor states, it does not preserve the additive structure of Eq. 1 in general, and we therefore do not assume that *Q* or *V* inherits the decomposition of *r*.

Building on this decomposed reward representation, we develop a probabilistic generative model of the joint policy *π*(*a* | *s*) by integrating a multi-agent reinforcement learning framework with maximumentropy value iteration [6, 25] (Section 4.3) and coordination strategies (Section 4.4). Using this generative model summarized in Fig. 1A, we can then infer the target reward function that best explains the observed behavior using the proposed inference algorithm (Section 4.5).

**Figure 1:**
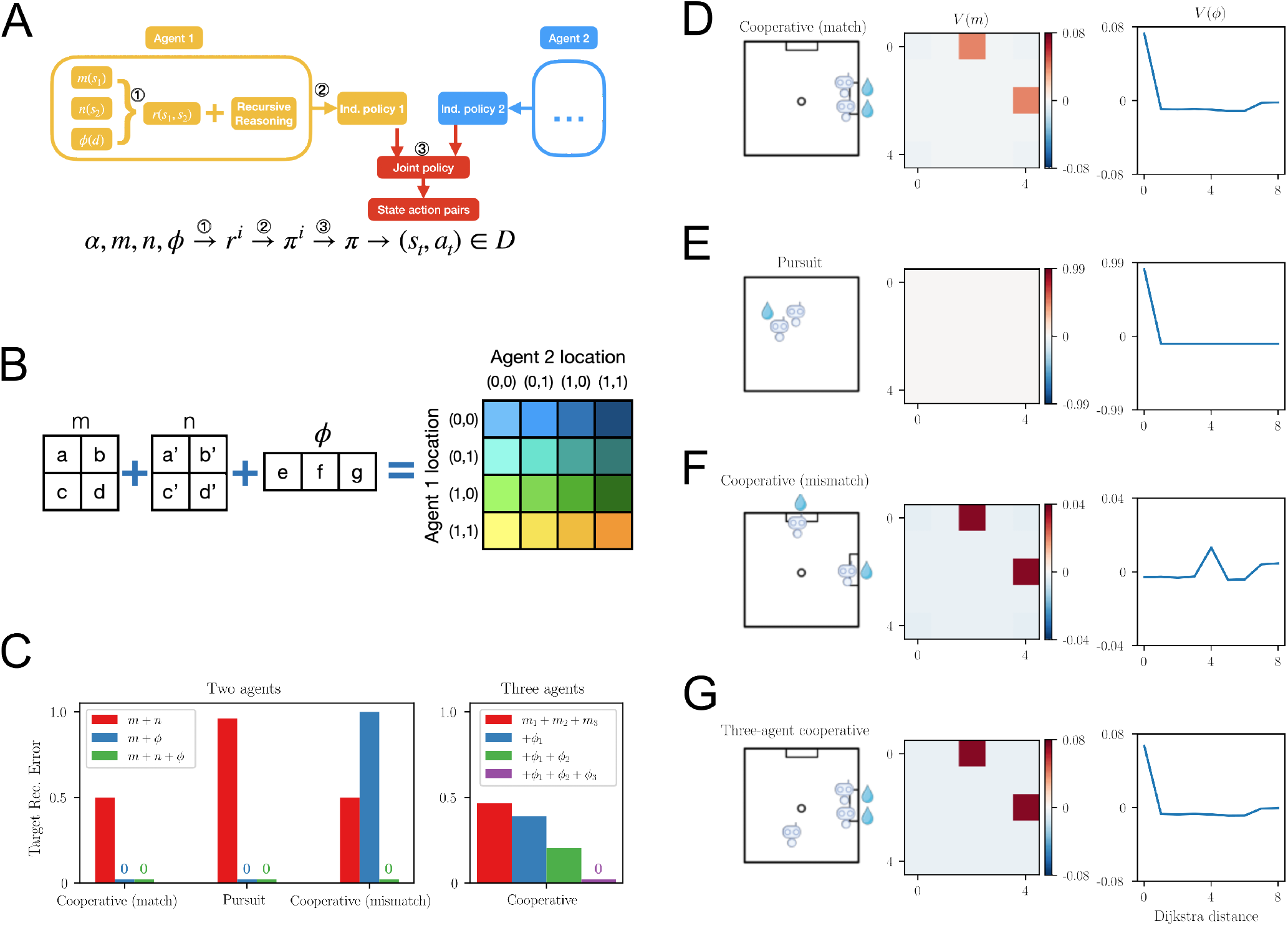
MAIRL and value decomposition for multiple social tasks. (A) Model overview. From observed joint state–action pairs (*s*_*t*_, *a*_*t*_), we infer each agent’s decomposed reward components *m, n, ϕ* based on a generative model of social decision making. The computation graph highlights the following stages: ① value decomposition (Eq. 1), ② policy generation with soft Q-learning (Section 4.3) and ③ coordination and cognitive levels (Section 4.4). (B) Value decomposition formulated as an overdetermined linear system. The actual reward function of joint locations *r*(*s*_1_, *s*_2_) is decomposed into summations of functions of each agent’s location *m*(*s*_1_) and *n*(*s*_2_) and their mutual distance *ϕ*(*d*). For two agents interact in a 2 × 2 grid world, 11 parameters *a, b, c, d, a*^*′*^, *b*^*′*^, *c*^*′*^, *d*^*′*^, *e, f, g* from three functions *m, n, ϕ* (left) are used to reconstruct the joint reward function *r* on the right with 16 free parameters. The design matrix is defined by the spatial relationships and grid layout (Fig. S1). (C) Target reconstruction error of the joint reward function from the ordinary least squares (OLS) solution of the value decomposition equation. Left, in all two-agent tasks under consideration, *m* + *n* + *ϕ* (green bars) suffices to achieve accurate reconstruction of target locations in the joint reward function. Right, in a three-agent cooperative task, incorporating each individual’s allocentric locations and their pairwise distances allows errorless target location reconstruction (purple bar). (D-G) OLS solutions of decomposed reward components *m*_*i*_, *ϕ*_*i*_ for different social tasks. Left, schematics for cooperative tasks in two-agent or three-agent cases; Middle: reward map of individual’s self allocentric location; Right: reward map of mutual distance.

### 2.2 Tractable reconstruction of joint reward functions via value decomposition

To illustrate how value decomposition recovers interpretable latent structure in social tasks, we consider a set of canonical multi-agent foraging paradigms, described in detail below. In each case, the task defines a sparse joint reward function over the combined state space. Although separability of the reward function *r* is not guaranteed in general, we show (Section 4.2) that for many social tasks the parameters *m, n*, and *ϕ* in Eq. 1 can be estimated using an ordinary least squares (OLS) solution (Fig. 1B and Fig. S1). Across these tasks, we evaluated reconstruction accuracy by measuring the proportion of incorrectly recovered target states as different components of Eq. 1 were included (Fig. 1C). The resulting reward maps and interaction terms provide a compact and interpretable description of the latent representations that guide behavior.

In the cooperative match task (Fig. 1D), two agents must reach the same goal location (north or east) simultaneously to obtain reward, resulting in two rewarded joint states. Value decomposition recovers this structure by combining individual reward maps that emphasize the goal locations with an interaction term *ϕ* that peaks at zero distance, reflecting the requirement for spatial proximity. Together, these components reconstruct the sparse joint reward function with zero error (Fig. 1C), and provide an interpretable representation of the task: agents are rewarded at both goal locations and for coordination with each other.

In the pursuit task (Fig. 1E), a reward is obtained whenever both agents occupy the same position, independent of their location in the arena. Accordingly, the interaction term alone captures the task structure, encoding reward as a function of inter-agent distance, while individual reward maps are not required. This demonstrates that value decomposition isolates the relevant latent variable, i.e., agent proximity, underlying the task.

In the cooperative mismatch task (Fig. 1F), agents must reach different goal locations simultaneously to obtain a reward. Here, value decomposition recovers individual reward maps that highlight the goal locations, together with an interaction term that encodes the preferred separation between agents. Notably, this interaction term peaks at a distance corresponding to the separation between the two reward zones, reflecting the specific coordination strategy required by the task.

In the three-agent cooperative task (Fig. 1G), reward is delivered when any two agents reach the same goal location simultaneously. Despite the large joint state space (25^3^ = 15,625 states), value decomposition recovers the reward structure using individual reward maps and pairwise interaction terms, greatly reducing the parameter space. These components capture both goal-directed behavior and pairwise coordination, demonstrating scalability to interactions with more than two agents.

Together, these examples demonstrate that value decomposition accurately recovers the structure of joint reward functions in a compact and interpretable form, revealing how social tasks can be represented in terms of individual goals and interaction constraints. This formulation enables efficient reconstruction with a tractable number of parameters across a range of multi-agent settings.

### 2.3 Validation of MAIRL in simulated cooperative foraging with heterogeneous strategies

Having established the accuracy of the OLS-based reconstruction, we next asked whether MAIRL can infer decomposed reward functions from simulated trajectories of agents with distinct reasoning strategies in the cooperative match task (Fig. 2A). Under independent action selection, the model should recover each agent’s reward function separately when the agents follow different decision strategies. Following Section 4.4, we simulated two agents with different reasoning policies: an egocentric, theory-of-mind level-0 (ToM-0) agent, which navigates to reward locations without considering its partner’s behavior, and a socially aware, theory-of-mind level-1 (ToM-1) agent, which assumes that its partner behaves as a ToM-0 agent and plans accordingly. Example simulated trajectories are shown in Fig. 2B and C. From behavioral observations alone, including trajectory length and port choice, these two strategies are difficult to distinguish.

**Figure 2:**
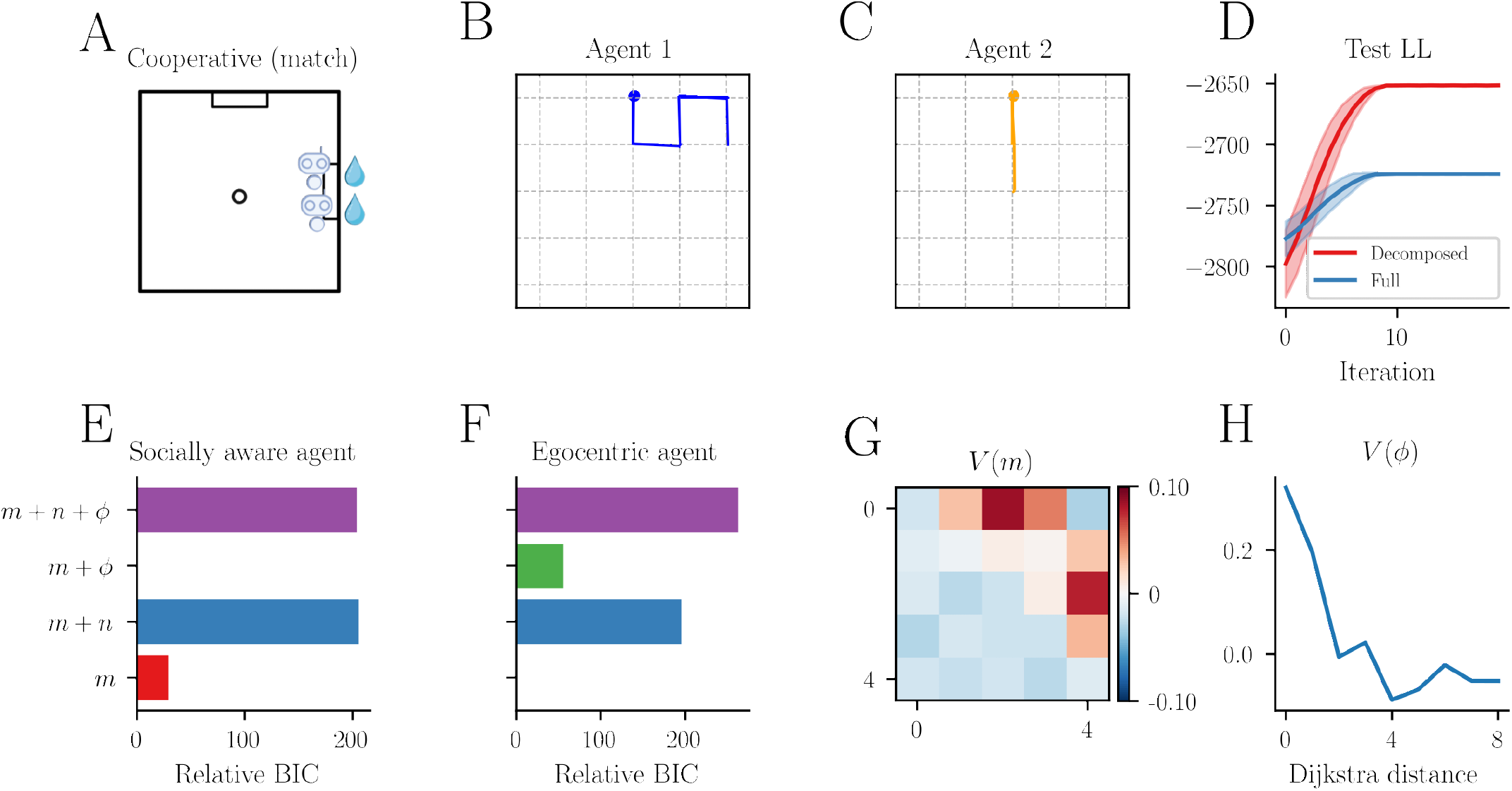
Recovery of reward functions in simulated two-agent cooperative match foraging. (A) Schematic of a simulated cooperative (match) foraging experiment in which reward is delivered only when both agents reach the target location simultaneously. (B–C) Example simulated trajectories of two agents in a 5×5 grid environment; solid dots indicate trajectory endpoints. (D) Test log-likelihood (LL) of the inference algorithm using either decomposed reward parameterization or the full joint reward function. Shaded regions denote variability across ten random initializations. The decomposed parameterization achieved higher LL, reflecting improved optimization due to reduced parameter dimensionality. (E-F) Relative Bayesian Information Criterion (BIC) of models with different structural components, fit to trajectories generated by (E) a considerate agent using a ToM-1 policy and (F) an egocentric solo-foraging agent. Each row corresponds to a model in which components of the value decomposition (Eq. 1) are added sequentially, enabling systematic hypothesis testing of behavioral strategies. Relative BIC values are normalized by subtracting the minimum BIC across model hypotheses. (G–H) Inferred reward maps for the ToM-1 agent, shown as functions of self-location and mutual distance, revealing strong weighting of target locations and proximal interaction states.

We then applied MAIRL separately to each agent’s trajectories to assess model fit and identify the reward components underlying their distinct strategies. First, behavioral fitting using the decomposed reward model achieved consistently higher log-likelihood (LL) than the full joint reward model across ten random initializations (Fig. 2D). This improvement is likely due to the reduced parameterization of the decomposed model, which makes the optimization less susceptible to local minima. Second, model comparison allowed us to assess which terms in Eq. 5 are necessary for capturing and distinguishing the agents’ underlying foraging strategies. The Bayesian Information Criterion (BIC) results of fitting the trajectories from both agents are shown in Fig. 2E and F. A BIC drop greater than 10 is generally considered strong evidence in favor of the lower-BIC model. For the socially aware agent (Fig. 2E), models composed of both self position and partner distance were preferred, whereas for the egocentric agent, the model containing self position only was sufficient. These selections align with the simulated ground-truth reasoning logic for each agent. The decomposed reward map inferred for the socially aware agent features elevated reward near the goal locations and in proximal states (Fig. 2G and H), mirroring the numerically decomposed reward structure for this task (Fig. 1D) and further validating the inference procedure. Together, these results show that MAIRL recovers agent-specific reward structures and supports principled hypothesis testing of behavioral strategies.

### 2.4 Benchmarking MAIRL against alternative multi-agent IRL methods

To ask whether value decomposition improves reward recovery relative to unconstrained multi-agent IRL, we compared MAIRL with MAMQL [26] and MA-AIRL [27] on the three simulated two-agent tasks (Coop. match, Coop. mismatch and Pursuit) defined in Section 4.7. These tasks use the reward rules of the examples in Fig. 1D–F, so that the ground-truth rewards are known. The methods were compared on the recovery of the rewarded joint states, the correspondence between recovered and true rewards, and the fidelity of the imitated policy (Section 4.7; Fig. 3).

**Figure 3:**
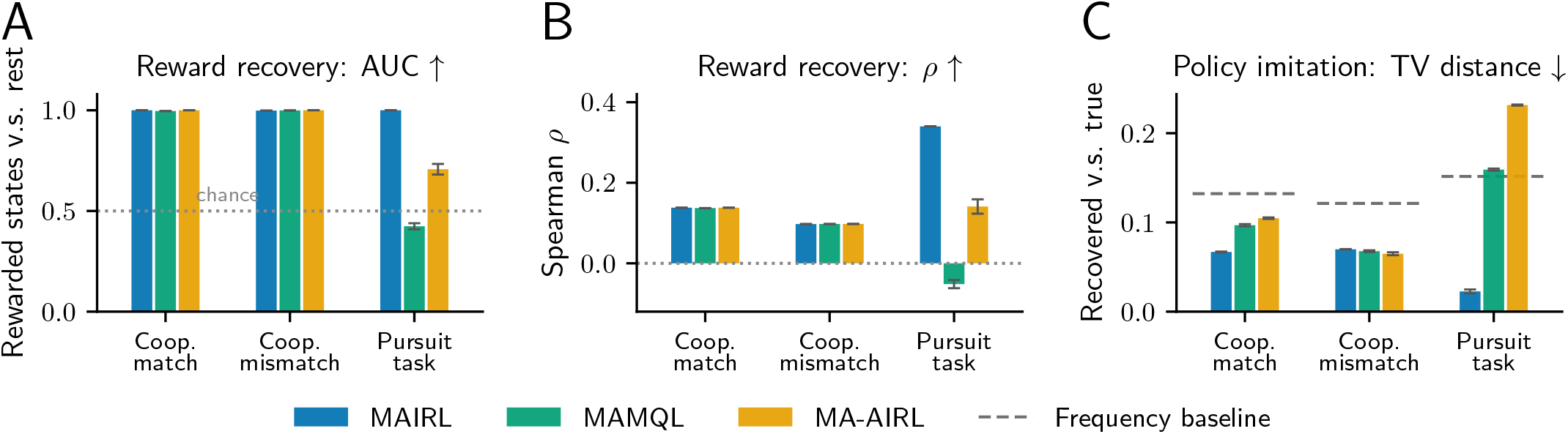
Benchmarking MAIRL against alternative multi-agent IRL methods in three simulated tasks. (A) Reward recovery scored as the ROC AUC with which the recovered reward ranks the true rewarded joint states above all remaining joint states (0.5, dotted line, chance). (B) Reward recovery scored as the Spearman rank correlation between the recovered and the true reward, evaluated over the joint states visited by the expert demonstrations. (C) Policy imitation scored as the mean total-variation (TV) distance between the recovered and the true focal-agent policy marginals (lower is better); dashed grey segments mark the empirical state–action frequency baseline, which provides a policy but no reward and therefore appears in (C) only. Bars are MAIRL (blue), MAMQL (green) and MA-AIRL (orange); each bar pools both focal agents and is the mean over five independently simulated datasets, each refitted with two optimization seeds. Error bars, s.e.m. across the five simulated datasets. Tasks differ in the reward rule: both agents on the same port (Coop. match, 4 rewarded joint states), the two agents on different ports (Coop. mismatch, 2 rewarded joint states), or both agents on the same cell (Pursuit task, 25 rewarded joint states).

MAIRL identified the rewarded joint states nearly perfectly in all three tasks (AUC = 0.998–1.000; Fig. 3A). The advantage of the decomposed representation was most evident in the Pursuit task, where the reward depends only on the relational structure between the two agents and the baselines degraded markedly (AUC = 0.424 for MAMQL and 0.707 for MA-AIRL). The rank correlation between recovered and true rewards was limited in the two cooperative tasks by their sparse reward structure, but remained highest for MAIRL in the Pursuit task (*ρ* = 0.340, versus 0.141 for MA-AIRL and ™0.052 for MAMQL; Fig. 3B). Policy imitation showed the same pattern: in terms of total-variation (TV) distance, MAIRL was closest to the ground-truth policy in the Coop. match and Pursuit tasks (0.067 and 0.023, versus 0.097 and 0.159 for MAMQL and 0.105 and 0.232 for MA-AIRL), whereas the three IRL methods performed similarly in Coop. mismatch (0.065–0.070; Fig. 3C). All methods outperformed the action-frequency baseline in the two cooperative tasks, but in the Pursuit task only MAIRL improved on it (0.023 versus 0.152). Together, these simulations show that the decomposed reward parameterization supports accurate identification of sparse rewarded states and accurate policy imitation across coordination rules. Imposing these additional structural constraints therefore does not harm performance in terms of either reward recovery or policy imitation.

### 2.5 Applications to experimental datasets

After validating our framework in simulations where ground truth is available, we applied the approach to the observed trajectories of two mice performing a cooperative foraging task [28] and tracked how their reward representations evolved during learning. This analysis revealed reward functions underlying distinct social roles and the unique dynamics in learning. Beyond cooperative settings, we further applied MAIRL to a non-cooperative “chicken” game in monkeys, demonstrating that the method generalizes across a range of social decision-making tasks.

#### 2.5.1 Infer reward functions in cooperating mice with distinct social roles

We first applied MAIRL to infer the reward representations underlying a cooperative match task performed by a pair of mice (Fig.4A). Similar to the simulated task, the two mice must cooperate by navigating to the same reward zones to obtain rewards [28]. The arena (45cm × 45cm) was discretized into a 10 × 10 grid, and animal movements were mapped onto this grid with nine possible actions (stay, up, down, left, right, and four diagonal movements). Example trajectory pairs are shown in Fig. 4B and C, where the animals selected the goal location in the east and coordinated their arrival.

**Figure 4:**
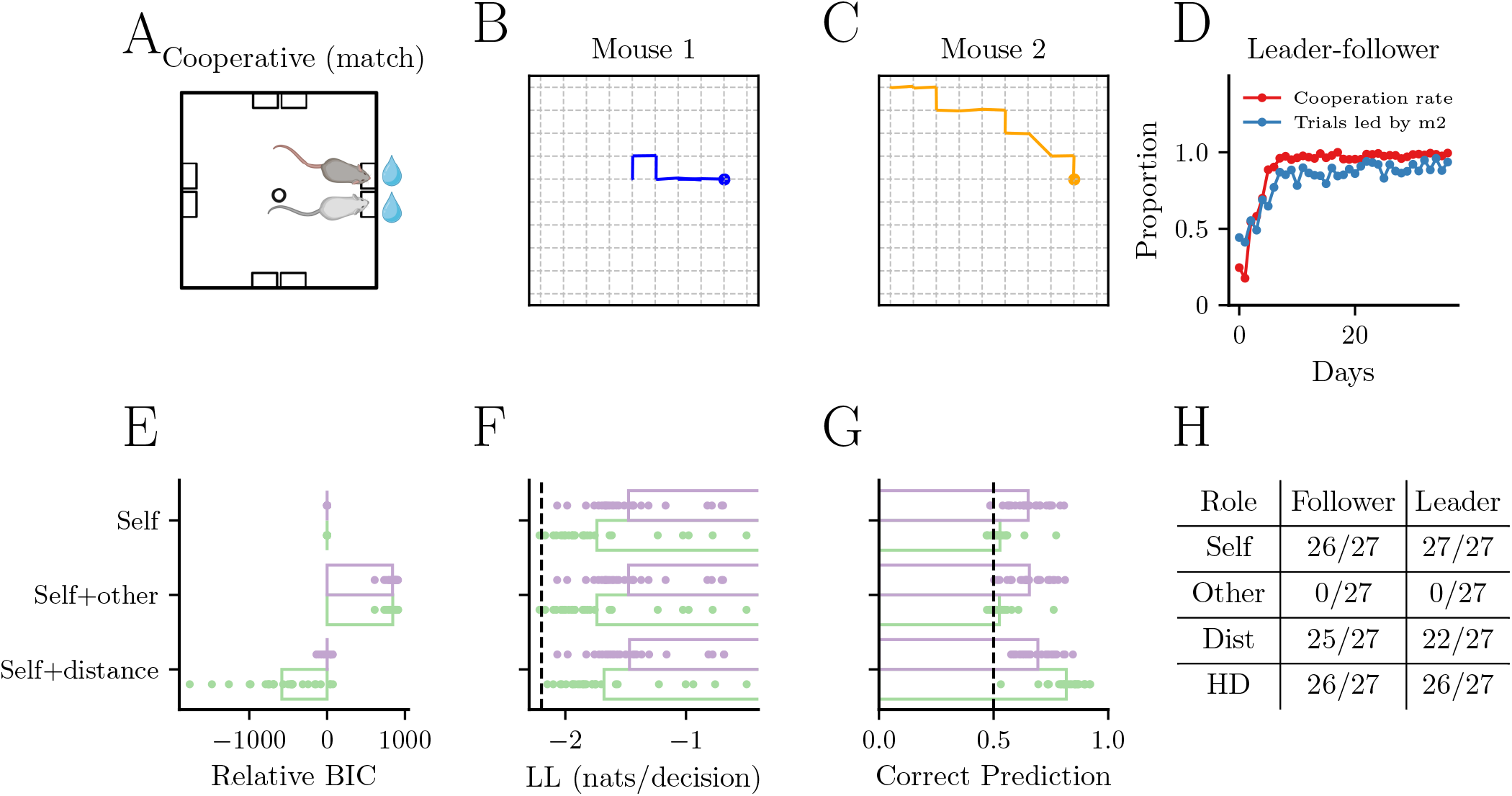
Hypothesis testing of decomposed value models reveals structured interactions and role-dependent strategies in mouse pairs. (A) Schematic of the cooperative foraging task in mice. Target locations are on the east and north. (B-C) Recorded trajectories from animal pairs navigating a 10 × 10 grid, reconstructed from empirical foraging data. (D) Proportion of cooperative trials and fraction of trials led by a single animal across training days. Over time, animals developed a consistent leader–follower strategy. (E) Relative BIC for 27 animal pairs, normalized to the minimal model containing only self-position value. Purple and green denote model fits for leader and follower animals, respectively. Each row corresponds to a fitting model in which different components in Eq. 1 are added sequentially. Adding allocentric other’s position did not improve model fit for either role, whereas including mutual distance significantly improved performance for both. (F) Same format as (E) but for log-likelihood per decision. The best-fitting pairs achieved values close to zero, indicating excellent behavioral prediction; the vertical dashed line indicates chance-level. (G) Prediction accuracy of the reward zone selected by animals, based on model-simulated trajectories. The dashed line indicates chance-level accuracy. (H) Number of animals (out of total) showing a significant improvement in model fit as each component was sequentially added. The self position model was first compared against a constant (uniform action) model; additional components were tested against the best-fitting simpler model using chi-square test. Almost all animals (26/27 followers, 27/27 leaders) showed significant self-position effects, and 26 of 27 follower and 26 of 27 leader animals showed significant head-direction modulation. Models involving head-direction (HD) are described in Fig. 7.

Mice adopt an emergent leader–follower strategy [28] to solve this task: the leader arrives first at the reward zone in most of the trials and has a disproportionate impact on follower’s choice. Using MAIRL, we sought to infer the reward functions underlying each animal’s decisions and to determine how these rewards relate to their social roles. To this end, we constructed a set of nested models to test whether each component of the decomposed reward model was necessary for explaining the behavior. We analyzed data from 27 well-trained animal pairs (Fig.4E-H). For most animals, models incorporating self-location and partner distance provided the best fit, whereas the partner’s allocentric location contributed minimally. Notably, partner distance improved model fit more for followers than for leaders. Overall, the best-fitting models significantly outperformed chance-level baselines (Fig. 4F) and accurately predicted final goal choices (Fig. 4G).

We then directly visualized the reward maps inferred from observed trajectories during cooperative task. Both leaders and followers (Fig. 5C, middle) showed higher reward at the goal locations. By contrast, followers showed significantly greater reward at proximal partner distance compared to leaders (Fig. 5C, right). Together, these results suggest that animals learn that successful coordination in this task depends primarily on maintaining appropriate mutual distance, with followers relying more heavily on partner distance compared to leaders.

**Figure 5:**
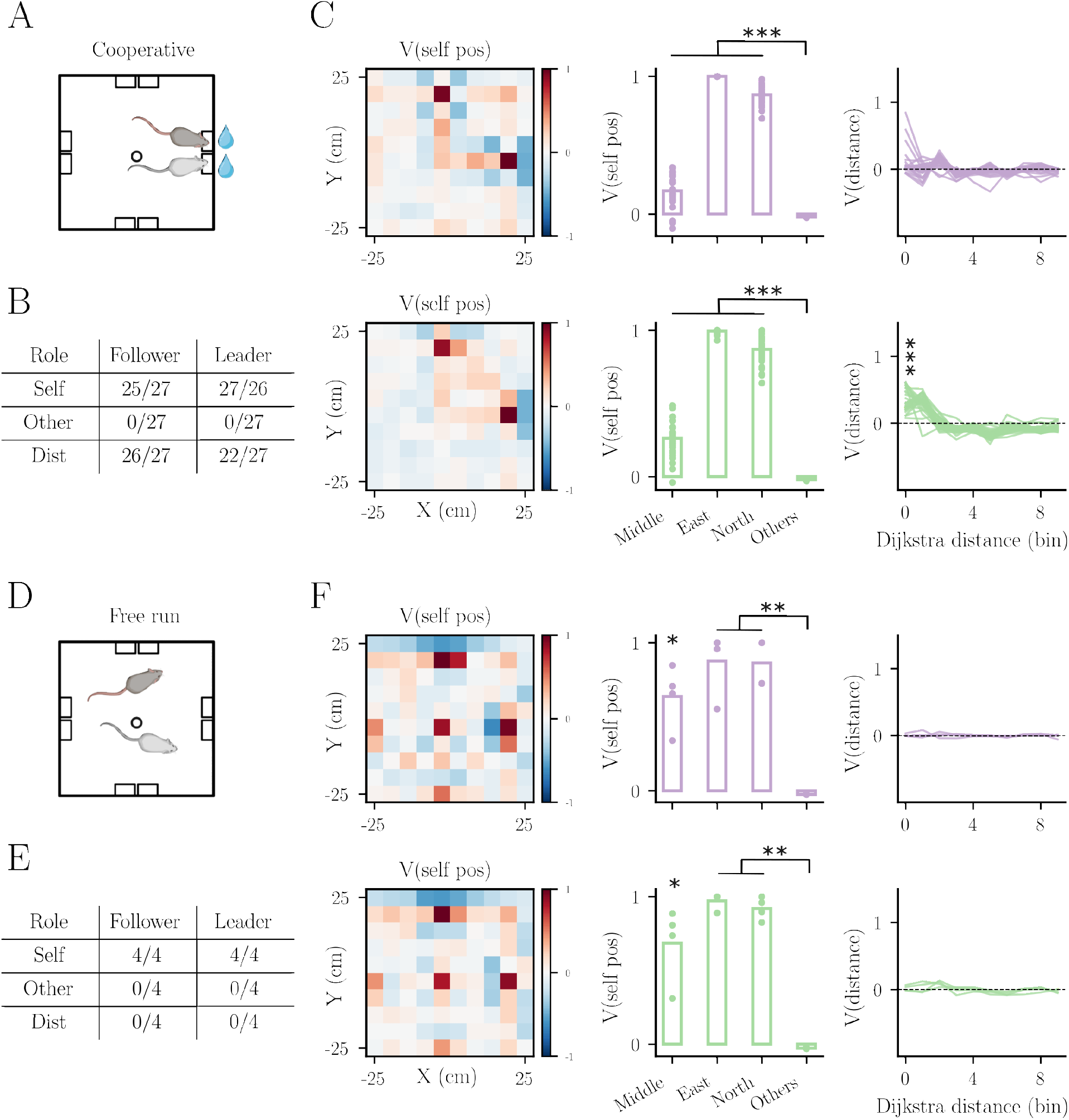
Reward maps inferred from observed trajectories during cooperative foraging and free-run sessions. *Cooperative task:* (A) Schematic of the cooperative task. (B) Number of animals showing significant improvement in model fit as components were sequentially added. (C) Inferred reward functions in correct trials for leaders (top row) and followers (bottom row). Left, example self-position reward map in allocentric space from one animal. Middle, reward assigned to the middle initiation spot and to the east and north reward ports, and the mean reward of all other locations across animals. Both leaders and followers show significantly higher reward at initiation and reward locations (paired t-test, \*\*\**P <* 10^*™*7^, *n* = 27). Right, inferred reward as a function of inter-agent distance. Followers show significantly higher reward at proximal partner locations than leaders (t-test at distance = 0, *P* = 6 × 10^*™*13^, *n* = 27). *Free-run:* (D–E) Same format as (A–B), but for free-run sessions. (F) Left, example self-position reward map. Middle, reward assigned to different locations across animals: middle locations show slightly elevated rewards compared to other positions (\**P <* 0.05), while target locations exhibit significantly higher rewards (\*\**P <* 0.01, *n* = 4). Right, inferred reward as a function of inter-agent distance is not significantly different from zero, consistent with the lack of improvement from including the distance component in Fig. 5E.

To determine whether MAIRL reliably reveals decision strategies across different behavioral contexts, we compared cooperative foraging to free-run sessions, when animals explore the arena out of the task context while still water-restricted (Fig. 5D). Model comparison (Fig. 5E) indicates that, in all four free-run pairs, both leaders and followers were best described by models in which decisions depended only on self-location. Although animals continued to show elevated reward at the goal locations (Fig.5F, middle)—likely because these sessions were usually recorded right before task sessions and the well-trained animals continued to visit goal locations in anticipation of potential reward—they no longer incorporated partner distance into their reward functions (Fig. 5F, right). This suggests that higher reward for partner proximity is selective during the cooperative task and absent when animals are not cued to coordinate.

Taken together, these results suggest that successful coordination in this task depends critically on the reward representation of partner distance, with followers relying more on partner proximity than leaders. Importantly, these reward maps are latent and not directly accessible from behavioral observations alone, but can be inferred through MAIRL. This demonstrates that the framework enables recovery of interpretable internal representations underlying social decision-making.

#### 2.5.2 Reveal the dynamics of reward components during learning

We next applied MAIRL to track reward components across learning, enabling a closer examination of the dynamics in reward assignment. The behavioral trajectories from the same 27 animal pairs were used. Because the reward components were estimated for each training session, we increased the discretization bin size and estimated rewards on a coarser 5× 5 grid.

Across learning, the reward component for partner distance progressively increased at close proximity over training in followers (Fig. 6A), while the corresponding representation in leaders showed little change (Fig. 6B). These results suggest that followers undergo greater reward updating during cooperative learning, selectively reinforcing proximity to their partners.

**Figure 6:**
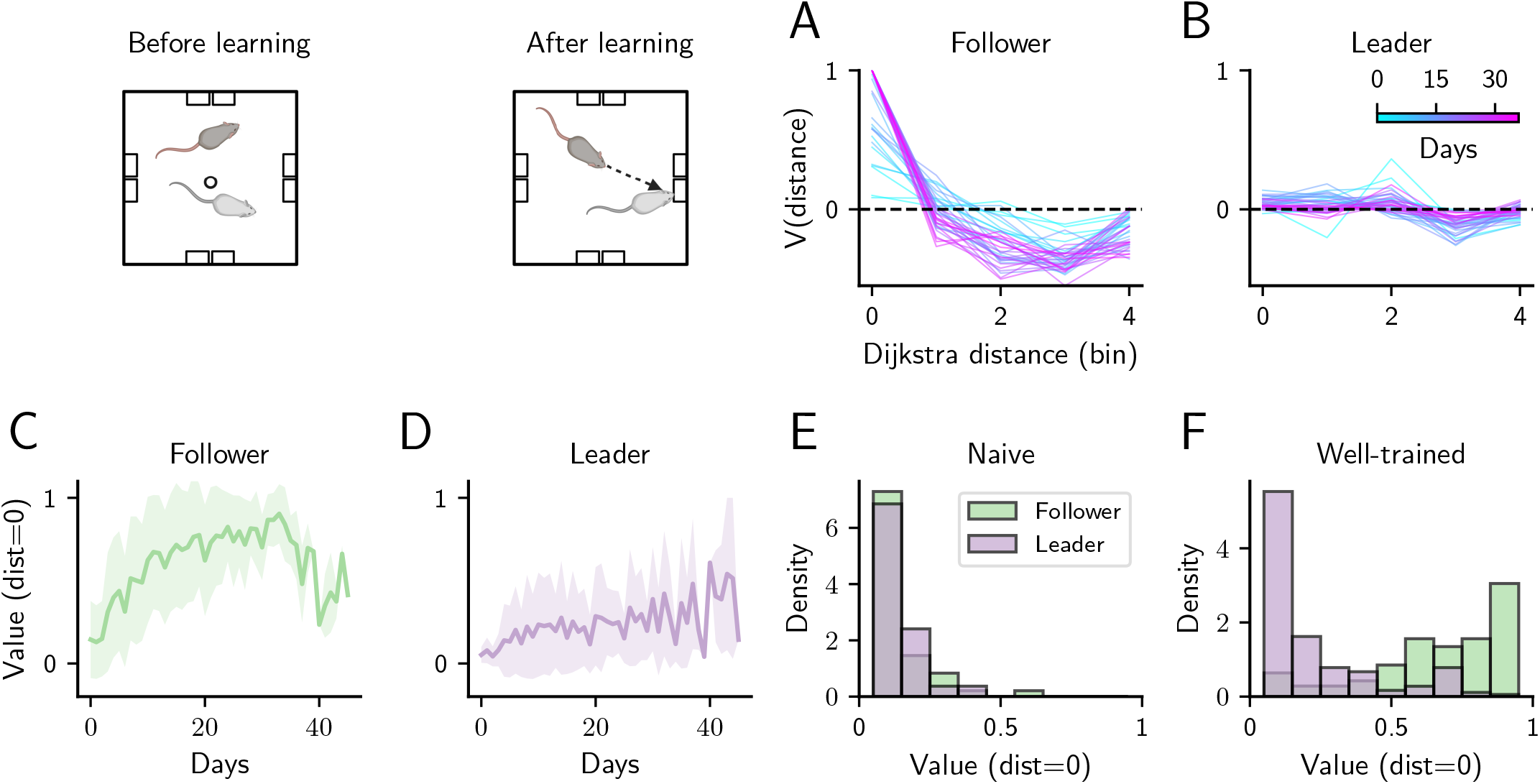
Dynamics of inferred reward components during learning. (A–B) Inferred reward as a function of inter-agent distance for a follower (A) and a leader (B), colored by training day. The follower progressively shows higher reward at proximal states, whereas the leader shows minimal change. (C–D) Reward assigned to the proximal state (distance = 0) over training days for followers (C) and leaders (D) (*n* = 27). Solid lines indicate mean rewards across 27 animals and shaded regions denote ±1 standard deviation. (E–F) Distribution of proximal-state rewards at the naive training stage (cooperation rate < 0.6; E) and at the well-trained stage (cooperation rate > 0.8; F). Histograms pool sessions with a significant distance component: naive, 85 follower and 90 leader sessions from 23 and 24 pairs, respectively; well-trained, 304 follower and 292 leader sessions from 27 pairs. For the comparison, rewards were averaged across sessions for each animal and followers versus leaders were compared with a paired t-test across pairs. Naive animals: followers and leaders do not differ (*n* = 23 pairs, *p* = 0.20). Well-trained animals: they clearly differ (*n* = 27 pairs, *p* = 5 × 10^*™*9^).

To quantify this effect at the population level, we pooled the rewards assigned to proximal partner distance across all 27 animal pairs over learning (Fig. 6C and D). During early learning, followers rapidly increased the reward of proximal partner distance, contributing to the increase in coordination success, whereas leaders showed slower and more limited adjustments. Consistent with this pattern, rewards for proximal partner distance, averaged across sessions for each animal, were indistinguishable between followers and leaders at the onset of training (Fig. 6E) but became significantly higher in followers after training (Fig. 6F). Together, these results reveal distinct learning strategies for followers and leaders, suggesting that role differentiation emerges through differences in reward updating. Importantly, MAIRL reveals these latent reward representations that are not directly observable from the behavior, providing a window into the internal computations underlying cooperative learning.

#### 2.5.3 Probing additional behavioral structure via conditioning variables

While the core MAIRL formulation captures social reward structure through a compact decomposition, it also enables hypothesis-driven incorporation of additional behavioral details without explicitly expanding the state space. Rather than introducing new state dimensions, we treat candidate variables as conditional modulators of existing reward components, allowing us to assess their contribution while preserving tractability. As a proof of concept, we examine whether relative head direction modulates the reward associated with inter-animal distance.

Relative head direction (*θ*) was defined as the angle of the partner from the instantaneous heading of oneself. We model *θ* not as an independent state variable, but as a conditioning variable that modulates the distance-dependent reward term:

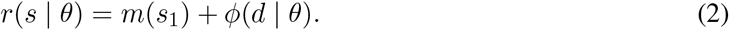

We discretized the continuous angle *θ* into three categories—front, left, and right—and fitted these conditioned distance reward maps to observed trajectories from the well-trained pairs (Fig. 7). Model comparison revealed that nearly all animals (26 of 27 followers and 26 of 27 leaders) showed significant improvement after introducing the additional parameters (Fig.4H). For these animals, conditioning on head direction reveals structured modulation of reward for partner distance (Fig. 7). Followers showed a stronger preference for proximal states than leaders when the partner was positioned directly ahead (front view: *p* = 10^*™*6^) or to the right (*p* = 5 × 10^*™*4^), whereas no significant role difference was observed when the partner was to the left (*p* = 0.10), suggesting that followers are more likely to approach and reduce distance when their partner is positioned ahead or to the right.

**Figure 7:**
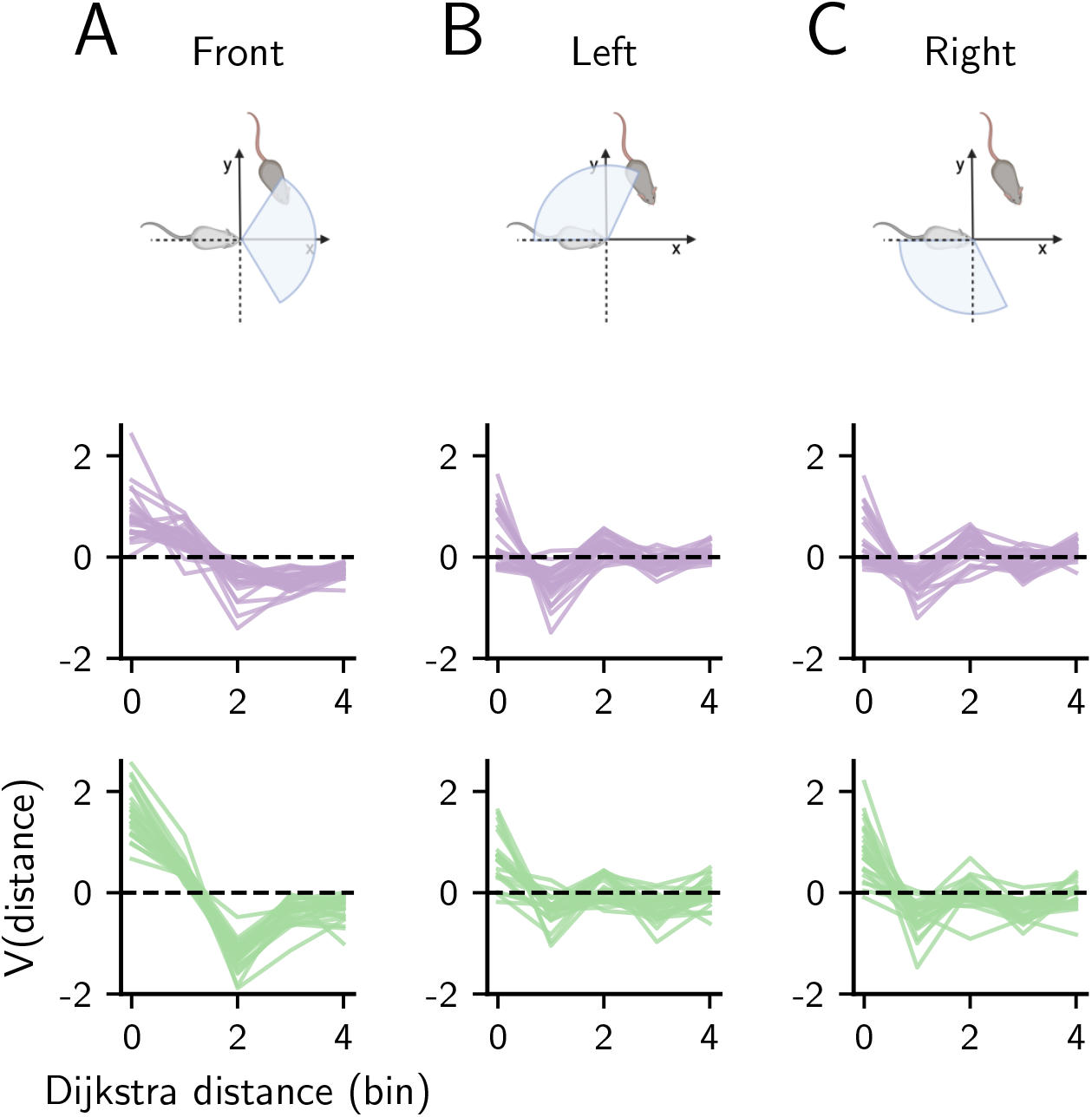
Distance-based reward functions conditioned on egocentric head directions. (A–C) Distance-based reward functions conditioned on the partner being in front (A), to the left (B), or to the right (C) of the focal animal. In each column, the top row shows the schematic of the partner-view condition, the middle row the leaders’ and the bottom row the followers’ reward functions. Each line is one animal for which the full model with self and distance terms fitted significantly better than a uniform-choice baseline and adding head direction significantly improved it; *n* = 25 leaders and 25 followers (25 of the 26 per role with significant head-direction modulation in Fig. 4H). Reward at distance 0 is above zero in every direction for both roles: • leaders, *p* = 2 × 10^*™*8^ (front), 10^*™*3^ (left), 10^*™*2^ (right); • followers, *p* = 2 × 10^*™*14^ (front), 5 × 10^*™*7^ (left), 6 × 10^*™*8^ (right); • followers versus leaders, front *p* = 10^*™*6^, left *p* = 0.10, right *p* = 5 × 10^*™*4^.

This analysis illustrates how MAIRL can flexibly test additional behavioral hypotheses without increasing the dimensionality of the state space. More broadly, it demonstrates that decomposed reward representations provide a structured framework for incorporating and evaluating candidate variables, enabling more precise and interpretable characterization of social behavior while preserving computational tractability.

#### 2.5.4 Infer mutual predictions in monkeys during a non-cooperative task

Extending beyond cooperative contexts, we used MAIRL to infer mutual predictions between monkeys in a non-cooperative “chicken” game. This non-cooperative game blends coordination and conflict and offers insights into players’ motivations and tendencies [29]. Two monkeys are presented with options: go Straight (S) or Yield (Y), resulting in four potential outcomes (Fig. S2A and B): (i) S-S leads to a crash with zero reward; (ii)/(iii) S–Y and Y–S yield a significant reward for the bold monkey and a small one for the yielding monkey; (iv) Y-Y results in cooperation with a moderate reward for both. Fig. S2B illustrates the outcome and payoff matrix from one monkey pair (trial number = 29,147), with decisions exhibiting roughly independent behavior, indicated by a mutual information measure of 0.01 bits.

This non-cooperative task is stateless with four possible actions in the joint action space. Two separate value functions, *V*_1_ and *V*_2_, need to be parameterized. The probability of agent 1’s decision could be written as:

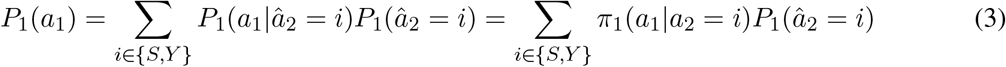

where *π*_1_(*a*_1_, *a*_2_) is a softmax transformation of *V*_1_(*a*_1_, *a*_2_) and denote *P*_1_(*â*_2_ = *S*) = *q*_1_. Inference details can be found in Section 4.10. The estimated *V*_*i*_ and *q*_*i*_ values (Fig. S2C-E) show that Monkey 1 estimates that its opponent has a lower probability of going straight. Consequently, Monkey 1 is bolder and chooses to go straight more often. Interestingly, the behavior of this monkey matches its higher social status as determined by a separate confrontation experiment [29].

Notably, the authors of these experiments [29] developed a “hybrid RL and strategic learning model” to determine one animal’s prediction about another, which is equivalent to a constrained MARL model with independent control, mutual logistic prediction and hard Q-learning update. Compared to this specialized model, our MAIRL model is more generally applicable to more complex predictions and state-dependent behaviors.

## 3 Discussion

We developed a multi-agent inverse reinforcement learning (MAIRL) framework that enables tractable and interpretable inference of latent reward representations in social decision-making. The key advance is a value decomposition formulation that represents joint reward functions as structured combinations of individual components and low-dimensional interactions, allowing recovery of high-dimensional social reward structure without estimating a separate reward parameter for every joint state. We show theoretically and numerically that this representation accurately reconstructs sparse joint reward functions while greatly reducing parameter complexity, with the corresponding value functions obtained by soft value iteration. Building on this, MAIRL enables direct inference of agents’ strategies from behavior: in simulated cooperative tasks, it distinguishes policies with different levels of social reasoning through model comparison. Applying MAIRL to mouse cooperative behavior, we uncover interpretable reward maps that reveal social role differentiation and distinct learning dynamics, with followers progressively increasing the reward assigned to socially relevant states. Importantly, the interaction terms in the decomposition are user-defined, allowing hypothesis-driven specification of candidate social features and direct testing of their contributions to behavior. MAIRL further supports the incorporation of additional behavioral features, such as head direction, as conditional modulators of reward representations without expanding the state space. Extending this framework to a non-cooperative “chicken” game in monkeys, we show that the inferred reward functions and mutual predictions capture strategic behavior and relate to social hierarchy, illustrating the generality of the approach across species and social paradigms.

The present formulation focuses on a minimal setting that isolates the core computational problem of inferring joint reward structure from behavior. We assume full observability of the environment and transition dynamics, allowing us to disentangle reward structure from uncertainty about others’ states. Extending the framework to partially observable settings would enable modeling how agents infer hidden states and beliefs during interaction. We further consider discrete state and action spaces without imposing a parametric form on the reward function, enabling flexible recovery of reward structure. Extensions to continuous domains using appropriate function basis would allow scaling to more complex environments while preserving this interpretability. Finally, the tasks studied here primarily engage low levels of recursive reasoning; applying the framework to settings that require deeper strategic inference would provide a more complete characterization of cognitive hierarchy in social behavior.

Together, these results establish MAIRL as a general framework for uncovering the computations underlying social interactions in biological and artificial systems. The framework naturally extends along several directions enabled by its decomposed structure. First, additional behavioral variables, such as motivation [30], movement kinetics [31], or vocal communication [32], can be incorporated to allow for systematic evaluation of their contribution to social behavior. Second, value decomposition enables scaling to larger groups by replacing exponential growth in the joint state space with growth that is linear in the number of agents for sparse interaction graphs (quadratic when all pairwise terms are included), making it possible to study structured interaction graphs and collective dynamics in larger social systems, such as schooling fish or social insects [5, 33, 34]. Third, applying MAIRL to tasks that require deeper strategic reasoning, such as the stag-hunt game [35], would enable quantitative inference of theory-of-mind capacities. These directions highlight that MAIRL provides a flexible and extensible foundation for investigating the computations governing complex social behavior.

Lastly, with the rapid development of large language model (LLM) agents, the proposed framework may also provide a useful perspective for studying interactions in artificial multi-agent systems. Recent advances have enabled multiple LLM-based agents to communicate, negotiate, and coordinate in shared environments, giving rise to increasingly complex collective behaviors [36]. However, the mechanisms governing these interactions often remain difficult to interpret, as policies and internal representations are distributed across high-dimensional neural networks. By explicitly modeling value functions, beliefs, and recursive reasoning, the MAIRL framework offers a principled way to infer latent objectives and predictive structures from observed social behaviors. This approach could help reveal how cooperative or competitive strategies emerge, how social roles differentiate within groups of agents, and how stable social conventions or equilibria arise over time. More broadly, applying such interpretable modeling tools to artificial agent societies may provide a quantitative bridge between studies of biological social cognition and the growing field of multi-agent artificial intelligence, enabling comparative analyses of learning dynamics, coordination strategies, and the emergence of collective behavior across natural and artificial systems.

## 4 Methods

### 4.1 Formulation of multi-agent inverse reinforcement learning (MAIRL)

Let’s consider a Markov Decision Process ^2^ described by ⟨*S, A, P, r*^1^, *r*^2^, *γ*⟩ where:

- *S* = *S*_1_ × *S*_2_ is the set of joint environmental states constructed by the cartesian product of individual state sets *S*_*i*_ with cardinality |*S*| = |*S*_1_| × |*S*_2_|
- *A* = *A*_1_ × *A*_2_ is the set of joint actions
- *P* = *P* (*s*^*′*^|*s, a*) : *S* × *A* → Δ(*S*) describes the state transition dynamics of the environment, where Δ is a probability measure over *S*.
- *r*^*i*^ : *S* × *A* × *S* → ℝ is the reward function that returns a scalar value to the *i*-th agent for a transition from *s* ∈ *S* to *s*^*′*^ ∈ *S*, taking joint action *a* ∈ *A*.
- *γ* ∈ [0, 1] is the discount factor for future steps.

In inverse reinforcement learning (IRL) [24][6][7], given {*P, S, A, γ*} and *N* samples of expert trajectories *D* = {*ζ*_1_, *ζ*_2_, ..*ζ*_*N*_ }, we aim to infer the unknown reward function *r*^1^, *r*^2^ such that *P* (*D*|*r*^1^, *r*^2^) is maximized. Each trajectory is composed of independent state-action pairs: 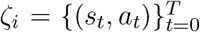 . As a consequence, the posterior probability of observing expert trajectories *ζ*_*i*_ can be calculated with the specific policy *π* derived from value functions (Eq. 4).

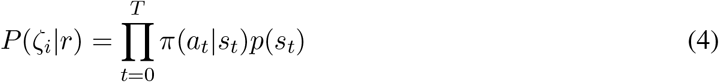

Therefore, at the core of this problem is to parameterize the *joint policy function π*(*a*| *s*) using a probabilistic modeling of social interactions based on MARL with recursive reasoning and value decomposition as highlighted in Fig. 1A.

### 4.2 Value decomposition

Inferring the reward function *r* : *S* → ℝ without assuming a parametric form requires estimating |*S*| parameters. In multi-agent settings, this becomes intractable, as the cardinality of grows exponentially with the number of agents. To address the issue, noting that *r* is usually sparse (see examples in Fig. 1D-G), we decompose the joint reward map into marginal maps and interactions maps as Eq. 5 where *S* ∈ *S*_*i*_ and *d* stands for a given interaction function between *s*_1_ and *s*_2_ (e.g., Dijkstra distance between agent 1’s location *s*_1_ and agent 2’s location *s*_2_ in a grid environment).

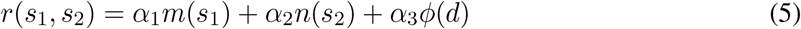

As a proof of principle, let’s first consider the case where the joint reward function *r*(*s*_1_, *s*_2_) : *S* → ℝ can be approximated by decomposing it into individual reward maps *m*(*s*_1_) : *S* _1_ →ℝ, *n*(*s*_2_) : *S* _2_ → ℝ and an interaction term *ϕ*(*d*) as a function of Dijkstra distance *d*(*s*_1_, *s*_2_). While this separability property is not always guaranteed, reconstruction error *r* ™ *m* ™ *n* ™ *ϕ* can be minimized through ordinary least square solution (OLS) of an overdetermined linear system.

Formally, we approximate the joint reward function *r*(*s*_1_, *s*_2_) by a separable decomposition of the form *m*(*s*_1_) + *n*(*s*_2_) + *ϕ*(*d*). This poses *O*(|*S*| ^2^) constrains to a system with *O*(2×|*S*|) parameters, constructing an overdetermined linear system. As depicted in Fig. S1, the reconstruction is expressed (Eq. 1) as *y* = *Xβ* where *y* is a flattened version of *r*(*s*_1_, *s*_2_), *β* is a concatenated vector of parameters in the *m, n, ϕ* and *X* is a fixed design matrix depending on the definition of the interaction term *d* and the grid environment. Since the number of constraints is much larger than the number of parameters, this is an overdetermined system. The ordinary least square (OLS) solution is expressed as 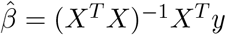 and the error vector evaluated as (*I* ™ *X*(*X*^*T*^ *X*)^*™*1^*X*^*T*^)*y*, denoted as *My*. The reconstruction error is the projection of vector *y* onto the null space of *M* . Since *M* and *y* both depend on the task setup and the choice of the interaction term, we may calculate the reconstruction error before performing the inference (Fig. 1C-G).

### 4.3 Policy parameterization

After parameterizing the reward function, we adopted a differentiable maximum entropy policy [6][25] to obtain a probabilistic mapping from reward functions to policies:

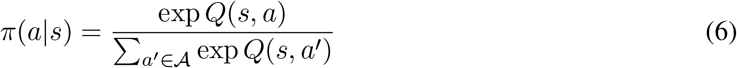

where *Q*(*s, a*) is a soft Q-function arising from performing soft value iteration:

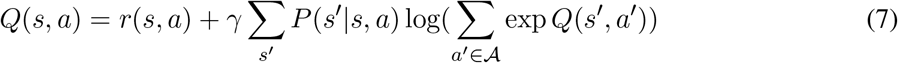

Note that a temperature term is not needed in the softmax function, as it will be absorbed into absolute value of *Q*, and subsequently *r*, which is the variable we aim to estimate from observations.

In the multi-agent scenario, the above generative model only holds for a cooperative task where two agents have the same reward function and act in a centralized way. In most realistic scenarios, however, agents have their own objectives and are not governed by a centralized controller. We therefore introduce separate reward functions, value functions, and policies *r*^*i*^, *Q*^*i*^ and *π*^*i*^(*a*^*i*^ | *s*) for each agent. If we assume that agents act independently and simultaneously, their joint policy could be expressed as Π_*i*_*π*^*i*^(*a*^*i*^ | *s*) . This will be the primary policy formulation to describe multiple agents in this work.

### 4.4 Coordination and theory-of-mind (ToM) levels

Another source of complexity in modeling multi-agent interactions arises from the need to account for coordination and recursive reasoning between agents. Each agent’s behavior depends not only on the environment but also on its beliefs about the strategies of others, introducing multiple levels of reasoning that must be modeled. For clarity, we focus on cooperative tasks in which agents share an identical reward function. We further assume that each agent’s state is fully observable to the others. The theory-of-mind (ToM) levels considered in this work, corresponding to different depths of recursive reasoning, are summarized in Table 1.

**Table 1:**
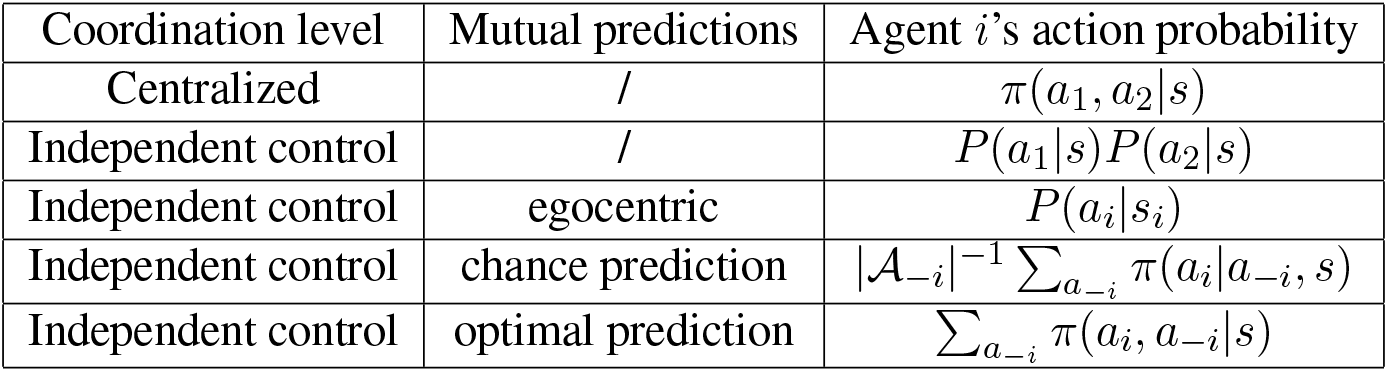
Generative models with different coordination levels.

| Coordination level | Mutual predictions | Agent *i*'s action probability |
|---|---|---|
| Centralized | / | π(a<sub>1</sub>, a<sub>2</sub>\|s) |
| Independent control | / | P(a<sub>1</sub>\|s)P(a<sub>2</sub>\|s) |
| Independent control | egocentric | P(a<sub>i</sub>\|s<sub>i</sub>) |
| Independent control | chance prediction | \|𝒜<sub>−i</sub>\|<sup>−1</sup> ∑<sub>a<sub>−i</sub></sub> π(a<sub>i</sub>\|a<sub>−i</sub>, s) |
| Independent control | optimal prediction | ∑<sub>a<sub>−i</sub></sub> π(a<sub>i</sub>, a<sub>−i</sub>\|s) |

In the fully coordinated scenario, agents behave as if controlled by a centralized planner, executing the optimal **joint action** defined by Eq. 6. In the following, *π* denotes this optimal joint policy, which may differ from the policies actually implemented by individual agents.

In a less coordinated scenario, agents independently execute their policy, which is often the case in many simultaneous action games. Mathematically, we have conditional independence of individual policies *P* (*a s*) = *P* (*a*_1_ *s*)*P* (*a*_2_ *s*). The question then becomes how each agent chooses its own action based solely on state information, without knowing the other agents’ action plans. The answer can be formulated through a probabilistic view of recursive reasoning, as described below:

For a ToM level-0 agent, which does not consider the action or state of other agents, its policy only depends on its own state information, i.e. *P* (*a*_*i*_ | *s*) = *P* (*a*_*i*_ | *s*_*i*_) and can be generated as soft value iteration of its own map function. For a level-1 or higher agent, it builds a predictive model of the other player and is expected to act optimally based on this prediction. Let’s denote agent 1’s prediction of agent 2’s policy as *â*_2_, based on which, we could express the policy of agent 1 as:

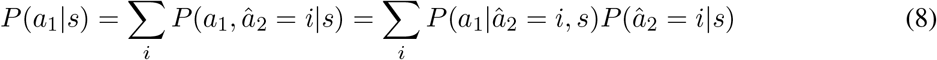

Since a level-1 agent acts optimally for the joint task based on its own predictions, the first term is essentially the conditional optimal policy *P* (*a*_1_ | *â*_2_, *s*) = *π*(*a*_1_ | *a*_2_, *s*). Thus, an agent’s policy is determined jointly by its prediction about its counterpart’s behavior and the underlying joint value function. The recursive level of this prediction defines the agent’s theory-of-mind (ToM) level. For example, if agent 1 assumes agent 2 to be a level-0 agent (regardless of whether agent 2 is or not), then agent 1 has a ToM level of one. If agent 2 assumes agent 1 is level-1 and forms its prediction about agent 1’s policy (i.e. *P* (*â*_1_ | *s*)) based on this assumption, agent 2 has a ToM level of two. Even higher levels of reasoning can be constructed using this recursive process. However, prior work suggests that human and animal behavior in strategic settings is typically well described by low levels of recursive reasoning [37–39]. Motivated by this and to maintain model tractability, we focus on low ToM levels that capture basic forms of social inference. Specifically, we consider two limiting parameterizations of mutual prediction under independent control: one in which agents make no informed predictions about their partners, and one in which agents assume their partners act according to the optimal marginal policy. These cases provide lower and upper bounds on coordination under this framework:

- If agent 1 has no information about agent 2’s policy, it generates chance level prediction (i.e. *P* (*â*_2_|*s*) = |*A*_2_|^*™*1^), then 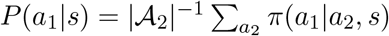 .
- If agent 1 predicts that agent 2 would act on the optimal marginal policy, it also acts on the marginal probability of the optimal joint policy. (i.e. *P* (*â*_2_ | *s*) = *π*(*a*_2_ | *s*)), then *P* (*a*_1_ | *s*) = Σ_*i*_ *P* (*a*_1_ | *a*_2_ = *i, s*)*π*(*a*_2_ = *i* | *s*) = *π*(*a*_1_ | *s*). Intuitively, when an agent correctly predicts that its partner follows the optimal policy, integrating over these predictions recovers its own marginal behavior under the optimal joint policy.

Most results in this paper, except Fig. S2, were inferred under the assumption of independent control with optimal prediction (last row in Table 1). These alternative assumptions about coordination and prediction can be evaluated through model comparison, as described later.

### 4.5 Inference procedure

After establishing a generative model based on the chosen hypothesis, our objective is to learn the map weights *α*_*k*_, individual value maps *m, n* and the interactions map *ϕ* that maximize the posterior of the observed expert trajectories *D* ={ (*s*_*t*_, *a*_*t*_) }.

The inference procedure is adapted from Ashwood et al. [7], noting that we have different components for the joint value function. We assume all parameters have a Gaussian prior with known variance and mean and individual maps have different prior variance compared to the interaction map. Mathematically, ***α*** ∼ *N*(**1**, Σ) where diag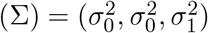, *m, n* ∼ *N*(**0**, Σ_*m*_), *ϕ* ∼ *N*(**0**, Σ_*ϕ*_). Note that we assume Σ_*m*_ and Σ_*ϕ*_ are diagonal matrices. Incorporating the map prior is equivalent to adding an L-2 regularizer with coefficients *λ*_1_ and *λ*_2_ to the map entries. The parameter optimization can be written as:

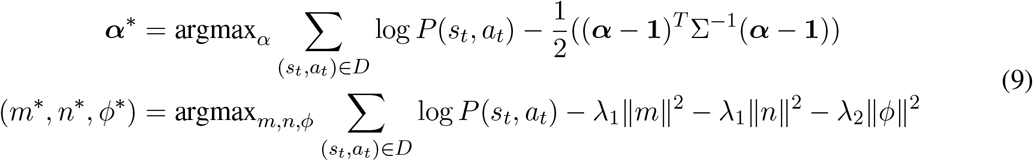

We performed coordinate ascent to iteratively update weights and maps while holding the other set of parameters fixed as illustrated in Alg. 1.

#### Algorithm 1

MAIRL with value decomposition

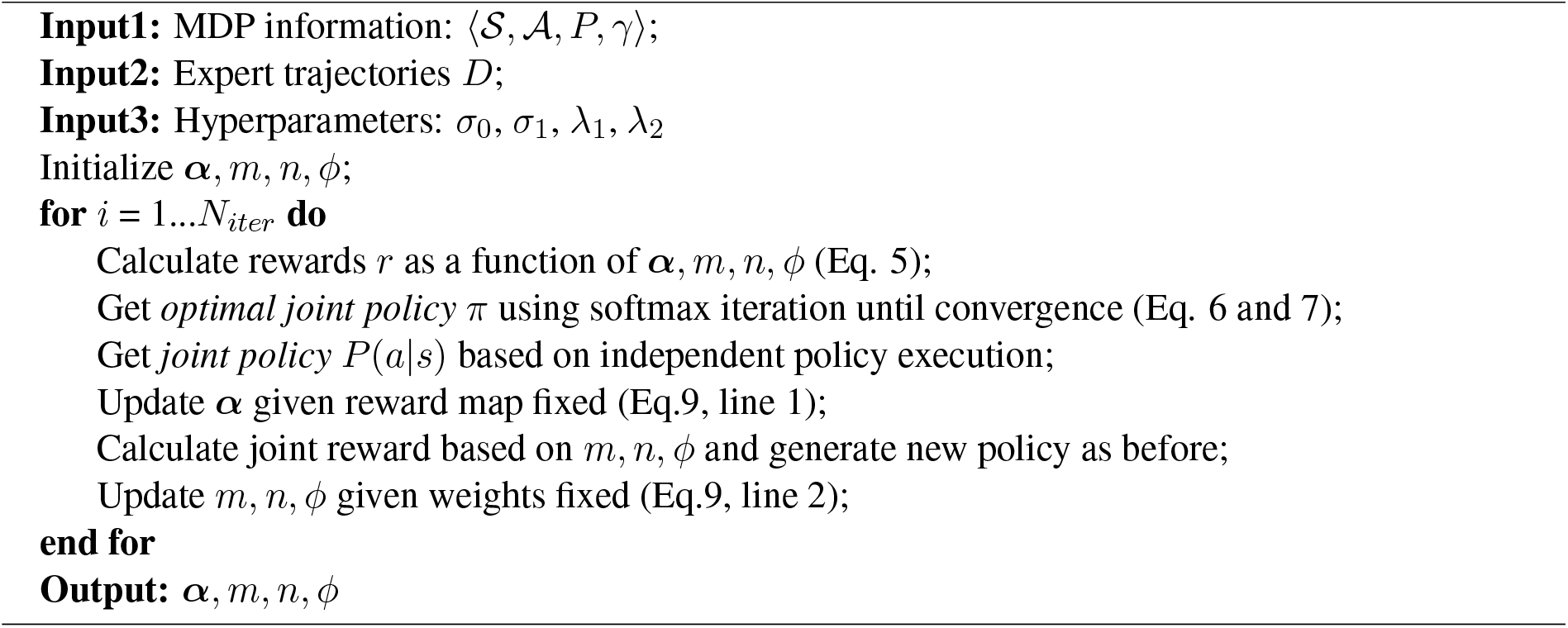

We used 80% trajectories for model fitting and the other 20% to calculate the test log-likelihood as model evidence. The preset hyperparameters include: the future discount factor *γ* = 0.9 and the covariance strength of map weights *σ*_0_ = *σ*_1_ = 0.01. In this way, we’re imposing a strong prior of the map weights to be around one, for a better comparison of the recovered maps.

Other hyperparameters were searched among the range provided in Table 2 in a centralized dataset inferred using centralized model assumptions. We picked *η*_*α*_ = 0.01, *η*_*map*_ = 0.005, *λ*_1_ = 5, *λ*_2_ = 1 based on training stability, model evidence and map interpretability.

**Table 2:**
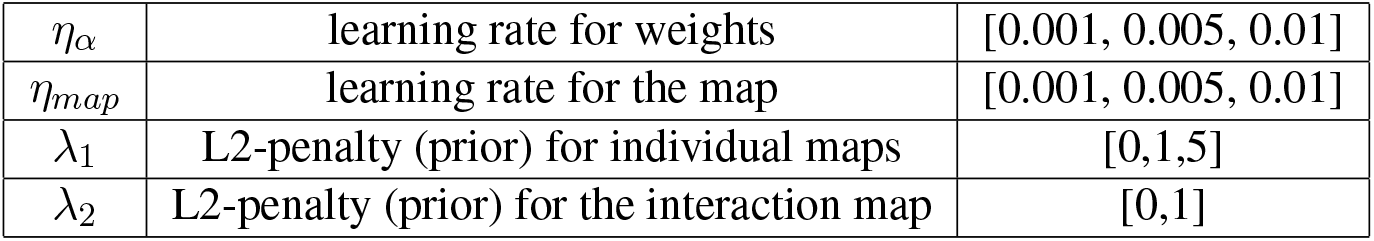
Range of hyperparameters tested.

|  |  |  |
|---|---|---|
| η<sub>α</sub> | learning rate for weights | [0.001, 0.005, 0.01] |
| η<sub>map</sub> | learning rate for the map | [0.001, 0.005, 0.01] |
| λ<sub>1</sub> | L2-penalty (prior) for individual maps | [0,1,5] |
| λ<sub>2</sub> | L2-penalty (prior) for the interaction map | [0,1] |

We initialized the weights to be one, the individual maps to be uniformly sampled from 0 to 1, and the location difference map to be 1*/*(*d* + 1) where *d* is the square Dijkstra distance between two agents. We used Adam optimizer with learning rate given above to iterate for 400 epochs. Most loss curves would plateau around 200 epochs. For 500 expert trajectories with approximately 30 decisions per trajectory, the inference process took around 5 minutes on a single CPU thread (Intel(R) Xeon(R) CPU E5-2687W 0 @ 3.10GHz).

### 4.6 Simulation of the cooperative foraging task

For the simulated trajectories shown in Fig. 2, the arena was discretized into a 5-by-5 grid with deterministic action transitions, resulting in 625 states and 25 actions in the two-agent MDP. Two terminal states are defined by the simultaneous presence of both agents at the same reward location either in the north or east. If an agent reaches the boundary of the arena, its position is reset to that of the previous time step. The reward strength at both goal locations was set to 2, as absolute reward magnitude does not affect the inferred reward structure. At the beginning of each trial, both agents are placed randomly in the arena. Agents are assumed to act under independent control. Agent 2 follows an egocentric (ToM level-0) policy, whereas agent 1 is modeled as a ToM level-1 agent that predicts agent 2’s behavior. In all simulations, 500 trials were generated, each with a maximum length of 200 steps. The discount factor *γ* was set to 0.9.

### 4.7 Benchmarking against alternative multi-agent IRL methods

#### Simulated tasks

Three two-agent gridworld tasks on a 5 × 5 arena were used. In Coop. match the arena contained four fixed ports, a four-port variant of the two-goal task in Fig. 1D, and a reward of 2 was paid only when both agents occupied the same port (4 rewarded joint states); in Coop. mismatch the two goal ports were at (0, 2) and (2, 4), and the reward was paid when the two agents occupied different ports simultaneously (2 rewarded joint states); in the Pursuit task the reward was paid when both agents occupied the same cell (25 rewarded joint states). Every trajectory had a fixed length of 30 steps and there was no absorbing state; the discount factor was 0.9. Expert demonstrations were generated from the ground-truth reward by soft value iteration (a centralized joint policy for Coop. match and Coop. mismatch, and an independent-control policy for the Pursuit task) and sampled as 2,000 trajectories per simulated dataset. Five datasets were simulated per task (seeds 401–405, 701–705 and 301–305), and each was split into 1,600 training and 400 held-out trajectories with a fixed split seed.

#### Recovery methods

MAIRL fits candidate reward decompositions by maximum entropy inverse reinforcement learning and selects among them by held-out BIC (Section 4.5; Fig. 2). For Coop. match the candidate set was self; self + other; self + dist; self + other + dist, where the interaction term is a nine-bin map of the inter-agent Manhattan distance, and self + other + dist was selected in all fits. Because the Coop. mismatch rule depends on which of the two ports each agent occupies, the interaction term was replaced by a two-dimensional map over the distances of the two agents to their nearest port (7 × 7 bins, jointly encoding how close each agent is to a port), and the candidate set was self; self + other; self + portpair; self + other + portpair; self + portpair was selected in all fits. For the Pursuit task the selected decomposition was self + dist in all fits. MAMQL [26] was fitted with the IQ-Learn objective (*χ*^2^ regularization, soft value *V* = log Σ exp *Q*, per-agent critics over the agent’s own five actions) for 5,000 updates, and MA-AIRL [27] with decentralized discriminators and PPO generators for 2,000 updates. Both baseline implementations used the same observation (one-hot own position concatenated with one-hot partner position), discount factor 0.9 and an additional critic weight penalty, and were fitted to both agents of each dataset. An empirical state-conditional action-frequency model, estimated from the training demonstrations, served as a policy baseline.

#### Evaluation

Every method was refitted with two optimization seeds per dataset, and both agents were scored as focal agents. Reward recovery was measured in two complementary ways. First, ROC AUC ranked the ground-truth rewarded joint states above all remaining joint states, which tests only whether the rewarded states are identified. Second, the Spearman rank correlation between recovered and ground-truth reward was computed over the joint states visited by the training demonstrations, which is invariant to monotone rescaling of the recovered reward. Policy imitation was measured as the mean total-variation distance between the recovered and the ground-truth focal-agent policy marginals over all joint states. For plotting, the two focal agents were averaged within each (dataset, optimization seed) cell, optimization seeds were averaged within each dataset, and the mean and s.e.m. were taken across the five simulated datasets (*n* = 5).

### 4.8 Animals, behavioral apparatus, and training

All procedures complied with the National Institutes of Health Guide for the Care and Use of Laboratory Animals and were approved by the Institutional Animal Care and Use Committee of the Icahn School of Medicine at Mount Sinai. Wild-type C57BL/6J female and male mice (2–6 months old; Jackson Laboratory) were housed under a reversed 12 h–12 h light–dark cycle (lights off at 07:00 and on at 19:00) in a temperature- and humidity-controlled environment (18–23 ^*°*^C, 40–60% humidity) with *ad libitum* access to food and water. Training began at 2–3 months of age, and all pairs were age-matched at the start of training. During training and testing, mice were water restricted but maintained at 80–90% of their initial body weight; each daily session lasted about 1 h, mice obtained 0.5–1.5 ml of water from the task, and supplemental water was given as needed.

Training took place in a 45 cm × 45 cm white-acrylic square arena. Four reward zones were centered on the arena walls, each with two adjacent water-delivery ports equipped with an infrared nose-poke detector and an LED indicating port availability. A central initiation detector registered trial onsets. Port signals were digitized (Arduino Nano; National Instruments USB-6001) and trial timing, LED signaling, and water delivery were controlled in LabVIEW; behavior was recorded at 30 frames per second with an overhead camera.

Mice progressed through four stages, one session per day. In stage 1, a single reward zone was separated from the rest of the arena by an acrylic barrier, and mice learned to poke the illuminated port (left or right, chosen at random) to obtain water. In stage 2, mice had access to the whole arena; a trial began when a mouse crossed the central initiation point, which illuminated both ports of one active reward zone, and a poke within 15 s produced water. In stage 3, two same-sex cage mates were introduced together; either mouse could initiate a trial, and both had to poke an illuminated port in the same zone within 15 s, with reward delivered only when the two pokes occurred simultaneously. In stage 4a, two of the four zones were illuminated after initiation, and the pair received water only if both mice chose the same zone (match rule). Mistakes (the two mice choosing different active zones, or either mouse poking an inactive zone) terminated the trial without reward and were followed by a 6–10 s timeout; omissions (no response within 15 s early in training or 8 s later) were excluded from the correct rate. Pairs reached criterion at ≥ 80% correct over three consecutive sessions and were excluded if criterion was not reached within 50 days.

To identify individuals, a small patch of back fur was shaved on one mouse of each pair. Nose, neck, and torso positions were tracked with SLEAP (v1.3.3) followed by Ensemble Kalman smoothing; the SLEAP model was trained on approximately 4,200 manually annotated frames from 10 pairs and applied to all sessions, and tracking was aligned to behavioral events using the LED onset/offset times. Leader and follower roles were assigned per session from the proportion of trials led by each mouse, tested against chance with a two-tailed binomial test (*α* = 0.01).

The analyses reported here used 27 well-trained pairs (54 animals). Five of these pairs were reported previously [28]; the remaining 22 pairs, together with the learning-stage sessions analyzed in this study, are new.

### 4.9 Pre-processing of mouse foraging trajectories

For inference using observed mouse foraging trajectories, each mouse was represented as a massless point anchored at its neck location. The arena was divided into a 10×10 grid with 5cm spatial bins (10cm bins in Fig. 6 and 7). An animal’s grid position was determined by assigning its actual location to the nearest grid point. To minimize artifacts introduced by discretizing the arena, we excluded time steps where both animals remained stationary. Each animal had nine possible actions: stay, up, down, left, right, up-left, up-right, down-left, and down-right. The objective was to estimate a value map that, given both animals’ current location, maximizes the probability of the observed action. The partner angle in Fig. 7 was defined as the angle of partner’s position relative to the agent’s own head direction. This angle was discretized into three bins: front (-60 to 60 degree), left (60 to 180 degree) and right (180 to 300 degree), such that *θ* ∈ {0, 1, 2 }. The discretized angle was included as an input to the model. To further limit model complexity, the decomposed joint reward function incorporates angle only through the interaction term. Analyses used sessions from the same 27 well-trained pairs (54 animals), drawn from two session sets. Behavioral model comparisons (Figs. 4, 5 and 7) used trajectories from well-trained sessions (cooperation rate ≥ 0.795; 437 sessions, of which 404 were retained after pre-processing, giving 47,335 correct trials), pre-processed as described above. Learning analyses (Fig. 6) additionally used all 745 sessions recorded from these pairs, including 117 training-stage sessions (cooperation rate *<* 0.6) and 191 intermediate sessions, so that the inferred reward could be tracked across training. We used the same held-out split across all model comparisons: 20% of each session’s trajectories (at least one) were held out, giving a median of 1,413 training and 356 held-out trajectories per pair.

### 4.10 Inference details for the chicken task

Unlike the previous two tasks, this non-cooperative task is stateless with four actions in the joint action space. Two separate value functions, *V*_1_ and *V*_2_, need to be parameterized. Value decomposition was not used due to limited number of parameters to estimate. The probability of agent 1’s decision could be written as:

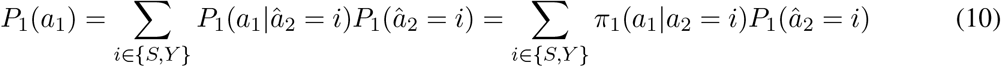

where *π*_1_(*a*_1_, *a*_2_) is a softmax transformation of *V*_1_(*a*_1_, *a*_2_) and denote *P*_1_(*â*_2_ = *S*) = *q*_1_. Therefore, *P*_*i*_(*a*_*i*_) is parameterized by five parameters each while the observed choice probability only offers two observation points. This creates an under-determined system. So we posed some constraints over the value function according to the task setup. We constrained *V*_1_ to be the transpose of *V*_2_ and *V*_1_(*S, Y*) *> V*_1_(*Y, Y*) *> V*_1_(*Y, S*) *> V*_1_(*S, S*) = 0 according to the task’s nature. In this way, we could have a faithful estimation of *q*_1_ and *q*_2_, which is usually the biggest ambiguity in these social interactions.

## 5 Data and code availability

The MAIRL code, documentation, and the simulated benchmark datasets are available at https://github.com/yschen13/ma_ir Of the mouse data analyzed here, five pairs have been published previously [28]; the remaining 22 pairs and the learning-stage sessions are available from the corresponding authors upon reasonable request. The monkey dataset was previously published [29]. All other data needed to evaluate the conclusions are present in the paper.

## 6 Author contributions

Y. Chen: Conceptualization, Methodology, Software, Formal analysis, Investigation, Visualization, Writing – original draft. Y. Cheng: Data curation. M. Kwak: Data curation. A. Radulescu: Conceptualization, Supervision, Writing – review and editing. H. Z. Wu: Conceptualization, Supervision, Funding acquisition, Writing – review and editing.

## 8 Acknowledgment

This work was supported by the Alfred P. Sloan Foundation, the Chan Zuckerberg Initiative, the Simons Foundation, the Esther A. & Joseph Klingenstein Fund (H.Z.W.) and the Friedman Brain Institute (H.Z.W. and A.R.); a Seaver Fellowship (Y. Cheng); a Swartz Postdoctoral Fellowship (Y. Chen).

## A Supplementary figures

**Figure S1:**
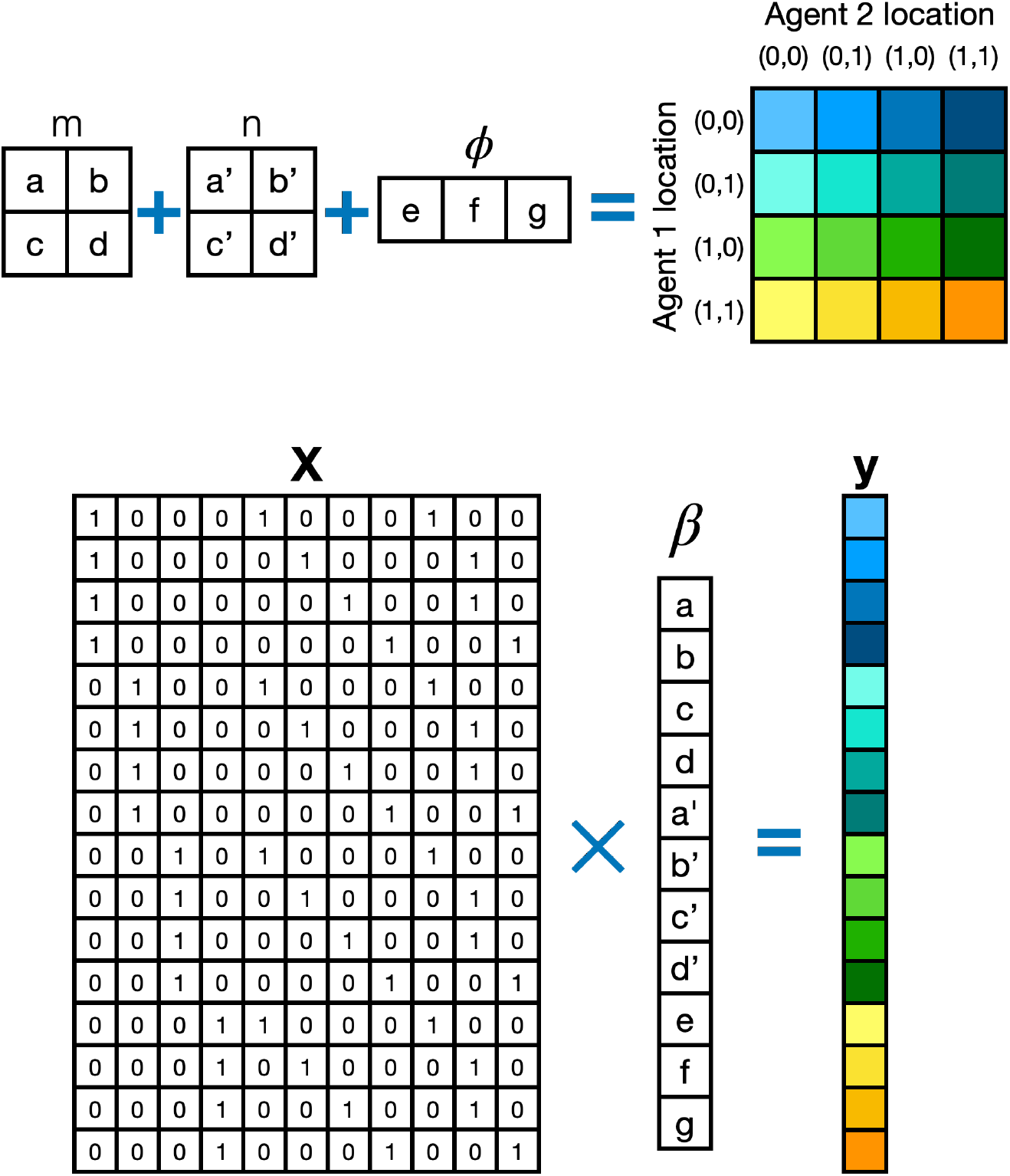
Value decomposition as an overdetermined linear system. Upper: for illustration, this task considers two agents interacting in a 2× 2 square arena, with the interaction value function *ϕ* defined over Dijkstra distance *d*. Eleven free parameters *a, b, c, d, a*^*′*^, *b*^*′*^, *c*^*′*^, *d*^*′*^, *e, f, g* from three functions *m, n, ϕ* (left) are used to reconstruct the joint reward function *r* with 16 fixed values depending on the task (right). The free parameters are defined as *a* := *m*(0, 0), *b* := *m*(0, 1), *c* := *m*(1, 0), *d* := *m*(1, 1), *a*^*′*^ := *n*(0, 0), *b*^*′*^ := *n*(0, 1), *c*^*′*^ := *n*(1, 0), *d*^*′*^ := *n*(1, 1), *e* := *ϕ*(*d* = 0), *f* := *ϕ*(*d* = 1), *g* := *ϕ*(*d* = 2). This constructs an overdetermined linear system with 11 unknowns and 16 constraints. Lower: to make this more intuitive, we could reshape the unknowns as a column vector *β* and reshape the fixed value functions as column vector *y*. Then design matrix *X* is then constructed based on the relationship between agents’ locations and the geometry of the grid environment. An optimal solution of *β* could be constructed as the ordinary least square (OLS) solution of this linear system: 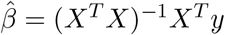 .

**Figure S2:**
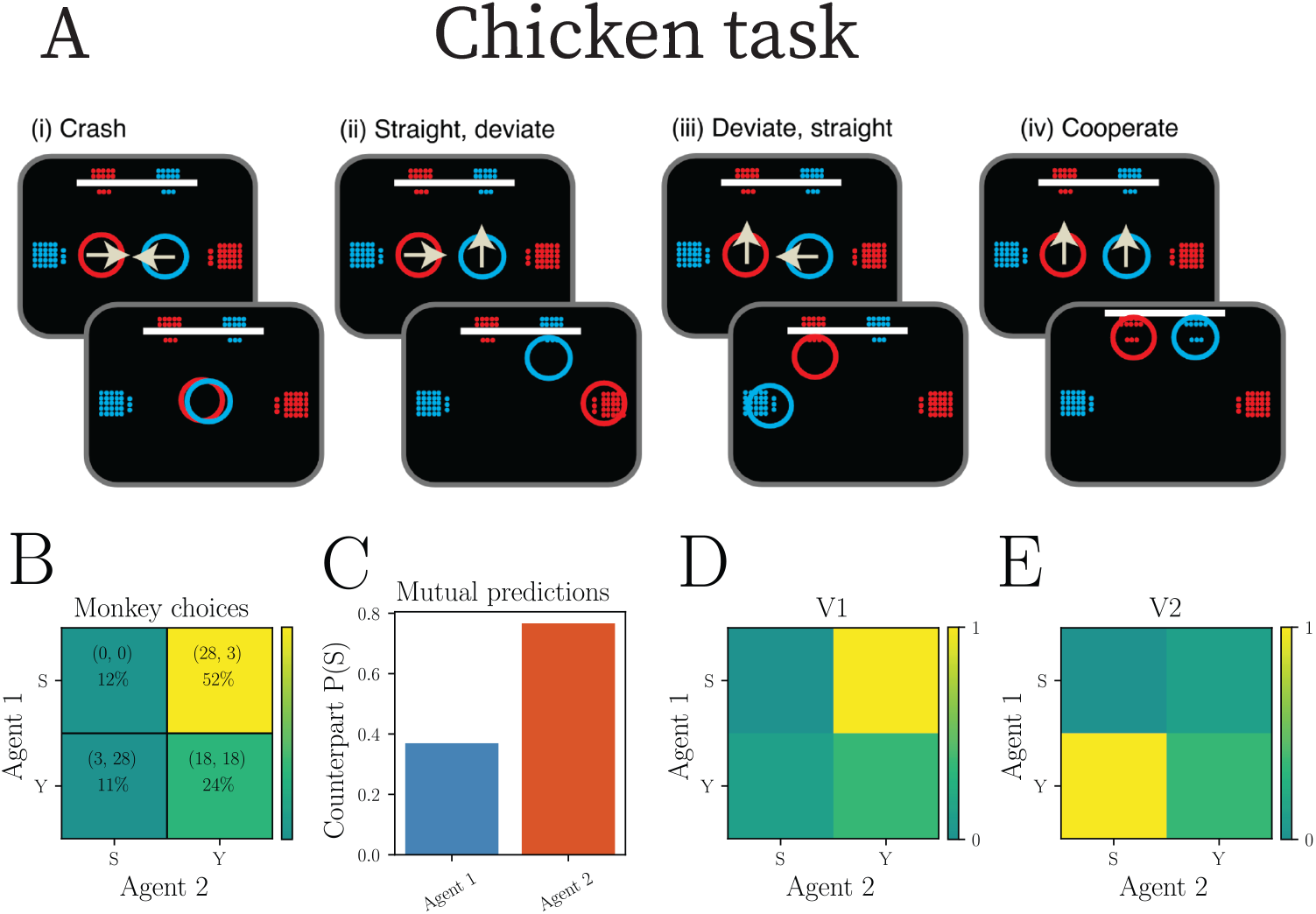
Monkey’s mutual predictions are related to their social hierarchy in a non-cooperative task. (A) Two monkeys (red and blue circles) choose to go Straight (S) or Yield (Y) in a non-cooperative chicken task. (adapted from [29]) (B) The choices of a pair of monkeys from the experiment (Fig. 2A in [29]). The value pairs in parenthesis indicate the pay-off matrix. (C) (D) (E) Estimated mutual predictions and value maps using a generative model of independent control with prediction. Mutual predictions are the probability of Monkey 2 choosing S as predicted by Monkey 1 and vice versa. Monkey 1 chooses to go straight more often because it is unlikely for Monkey 2 to go straight. This strategy matches their social hierarchy where Monkey 1 has a higher rank than Monkey 2.

## Footnotes

1 For clarity, we present the two-agent case; the formulation extends naturally to more agents, at the cost of exponential computational growth.

2 For simplicity, we provide equations for two agents. These derivations naturally extend to many agents albeit with exponential growth of required computation.

